# Species-specific small models for cell type classification approach the performance of large single cell foundation models

**DOI:** 10.64898/2026.03.16.711196

**Authors:** Gita Mahmoudabadi, Lakshmi Krishnan, Tejaswini Ganapathi, James Pearce, Steve R. Quake, Theofanis Karaletsos

## Abstract

Accurate cross-species cell type classification remains a key evaluation task in single-cell transcriptomics. Recent foundation models trained on millions of single-cell profiles demonstrate great in-distribution and out-of-distribution performance on this task, but their large parameter counts and substantial computational costs limit accessibility and interpretability. Here, we introduce CytoType, a simple and interpretable model for cell type classification that leverages pre-trained ESM-2 protein embeddings of protein-coding transcripts. By learning linear, cell-type-specific weights over transcript embeddings, without relying on gene count information, CytoType achieves F1 scores comparable to or exceeding those of large-scale transformer-based models. We further developed ESM-Cell Embedding (ESM-CE), an even simpler variant that only averages ESM-2 embeddings across expressed genes, which also performs competitively against foundation models. Both CytoType and ESM-CE are trained on species-specific data, maintaining high accuracy when classifying cell types with orders of magnitude fewer parameters compared to larger foundation models. For example, for human tissues, the average performance gap between CytoType and the best foundation model was 0.053 F1 points while CytoType uses 10,000x fewer trainable parameters. Additionally, we quantified the contribution of ESM-2 embeddings to cell type classification tasks and demonstrated a three fold reduction in the performance gap between CytoType and the best foundation model for nine species. Finally, we show that CytoType’s learned gene weights are biologically interpretable.

## 1 Introduction

Single-cell RNA sequencing (scRNA-seq) has revolutionized our understanding of tissues and cell types by profiling gene expression at the resolution of individual cells, but accurate cell type classification remains an important task for various downstream tasks and novel biological discoveries. Early cell type classification methods roughly fell into two major categories: marker-dependent approaches that rely on expert-curated gene sets, and profile-based classifiers that map cells to reference atlases (e.g. SingleR (Aran et al., 2019), scmap (Kiselev et al., 2018), and CHETAH (de Kanter et al., 2019)). However, these methods often struggle to generalize to different species and can be laborious as some involve manual curation of marker genes.

Inspired by the success of transformer foundation models in natural language processing, several groups have developed large single-cell transcriptomic foundation models such as scBERT (Yang et al., 2022), scGPT (Cui et al., 2024), Geneformer (Theodoris et al., 2023), scFoundation (Hao et al., 2024), UCE (Rosen et al., 2023), and TranscriptFormer (Pearce et al., 2025). While these foundation models offer the potential for many downstream tasks besides cell type classification, their multi-million-cell pre-training and large model sizes impose steep computational costs if the intended task is limited to cell type classification. Hence, we aimed to develop a simple yet effective linear baseline for cell type classification using transcriptomic data and pre-trained ESM-2 protein embeddings for protein-coding transcripts. ESM-2 is a transformer-based protein language model trained on unlabeled protein sequences with a masked language modeling objective. As a result ESM-2 implicitly learns proteins’ structural and functional signals (e.g. residue-residue interactions), whose embeddings can be used as a pan-species vocabulary for downstream modeling tasks.

Here, we propose CytoType, a lightweight and interpretable model that does not require marker gene selection and strikes a balance between simple classifiers and computationally-expensive foundation models. By embedding each transcript sequence with ESM-2 (Lin et al., 2023), and then learning cell-type-specific linear weights over these embeddings (Figure 1, Materials and Methods), CytoType achieves F1 scores competitive with many transformer-based approaches (Figures 2–4), while avoiding massive pre-training computational costs (Figure 5). The transcribed genes are selected via three methods, leading to three variants of CytoType, a) CytoType-ESM-Max, which uses genes sorted by maximum expression, b) CytoType-ESM-HVG, which uses highly variable genes and c) CytoType-Random-Max, which uses random gene embeddings for the maximally expressed genes, which serves as a basic baseline. All three CytoType variants use up to 2048 expressed genes.

**Figure 1:**
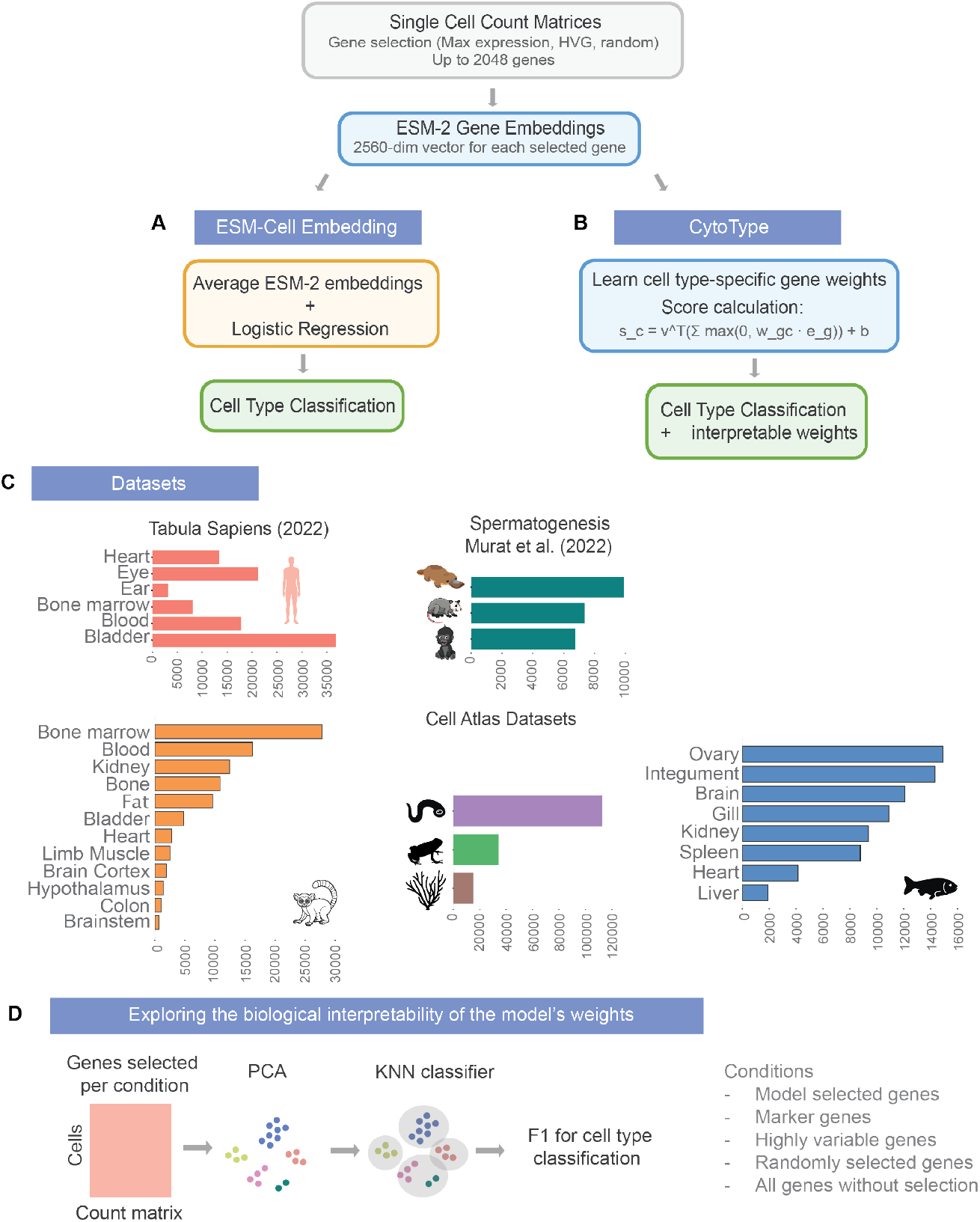
Schematic overview of models (A, B), data used during training and evaluation (C), and (D) analysis of model weight interpretability. A) ESM-Cell Embedding (ESM-CE). For each cell, the ESM-2 embeddings of top 2048 most expressed genes are retrieved and averaged to produce a single cell-level embedding. A logistic regression classifier predicts cell type from this averaged embedding. No gene-specific parameters are learned. B) CytoType. Expressed genes are first filtered by one of three gene-selection strategies (Max, HVG, or Random). ESM-2 embeddings are retrieved for each selected gene. The model learns a matrix of linear weights assigning gene importance for each cell type. These weights, applied to the corresponding ESM-2 embeddings, produce cell-type scores used for classification. Learned gene weights enable biological interpretability. C) Single-cell datasets used for model training and evaluation. The datasets span Tabula Sapiens, Spermatogenesis including testes tissue from three species (platypus, gorilla, and opossum) and Cell Atlas datasets containing Mouse Lemur (multi-tissue), Zebrafish (multi-tissue), Lamprey (brain), Frog (whole body) and Coral (whole body) datasets. D) Comparison of cell-type classification F1 scores based on va_1_r_7_ious gene-selection conditions to analyze the interpretability of model weights.

**Figure 2:**
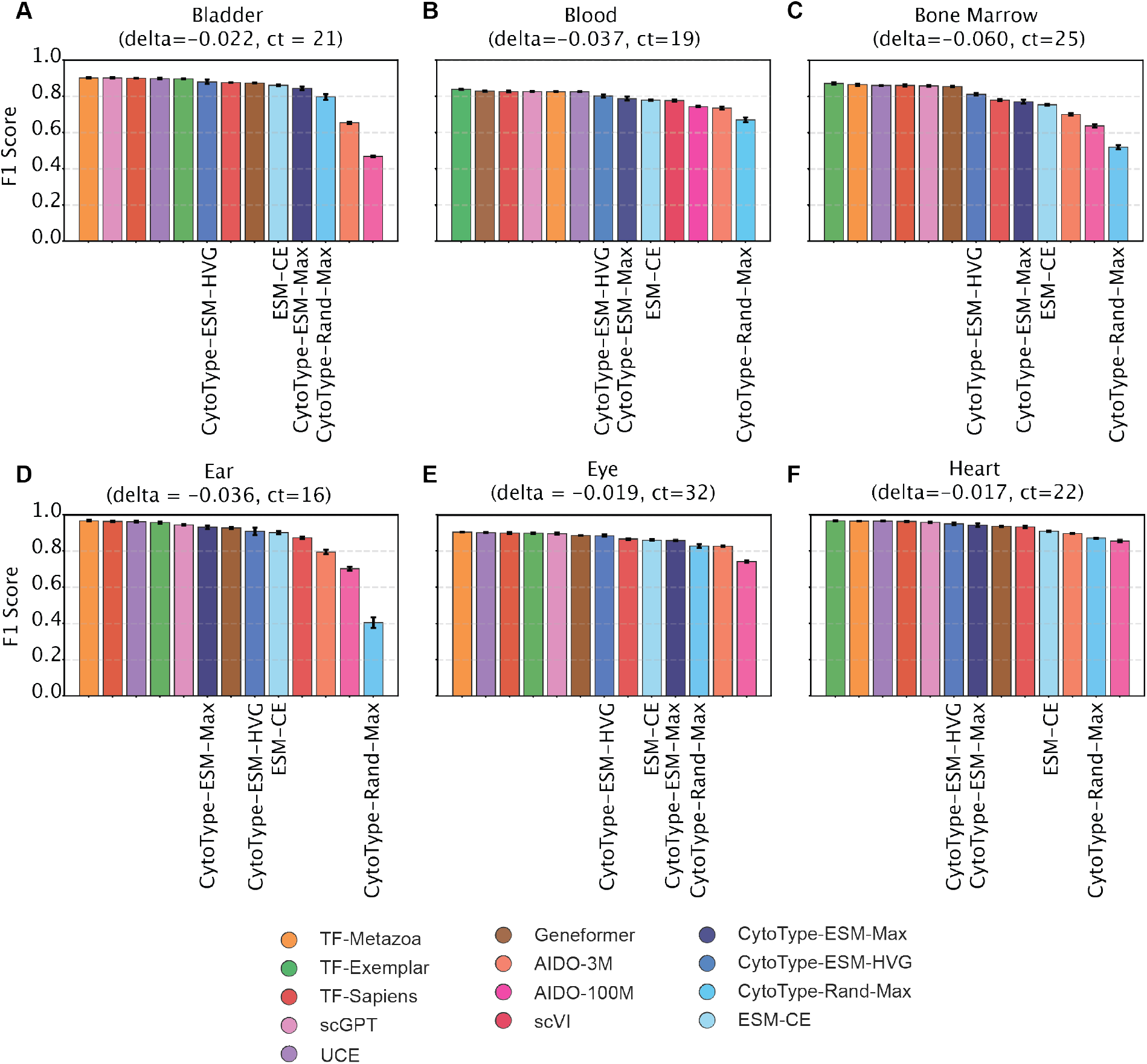
Cell type classification benchmarks for human tissues. A-F) Macro F1 scores benchmarking CytoType variants and ESM-CE against several single-cell foundation models. The difference between the best performing light-weight model and the best performing foundation model is denoted as delta. The number of cell types in each dataset is represented by “ct”. The models are sorted by their mean F1 score, from highest to lowest.

In addition to CytoType, we developed an even simpler model called ESM-Cell Embedding (ESM-CE) that does not learn cell-type specific gene weights. ESM-CE uses the average of ESM-2 embeddings for protein-coding transcripts that are expressed in each cell as an input to a Logistic Regression classifier (Figure 1, Materials and Methods). Although this model does not provide much interpretability, it performs quite competitively not only against CytoType, but also against other models (Figures 2–5). We further summarize the performances of different models across nine species as a function of the model’s computational complexity as measured by the number of trainable parameters (Figure 6). In the sections that follow, we describe both models in more detail, discussing their benchmarking results across tens of tissues and nine species. We also demonstrate how CytoType’s gene-specific weights carry information about which genes are specific to each cell type, and thus offer interpretability (Figure 7). Overall, we show that these simple models could achieve comparable performance to foundation models for cell type classification even though they require 4-5 orders of magnitude fewer trainable parameters and do not include absolute or relative transcript counts during training.

## 2 Results

### 2.1 CytoType and ESM-CE performance comparison against large-scale foundation models

We compared the performance of CytoType variants and ESM-CE to several single cell foundation models including TranscriptFormer (TF), which represents a family of large-scale single-cell foundation models trained on up to 112 million cells spanning more than 1.5 billion years of evolutionary history with three variants: 1) TF-Metazoa trained on 112 million cells from all twelve species, including six vertebrate species (human, mouse, rabbit, chicken, frog, zebrafish) four invertebrate species (sea urchin, C. elegans, fruit flu, and fresh water sponge), Saccharomyces cerevisiae (a fungus) and Plasmodium falciparum (a protist). Its total number of trainable parameters is 444 million. 2) TF-Exemplar trained on 110 million cells from human (H. sapiens) and four model organisms: mouse, zebrafish, fruit fly, and C. elegans, with 542 million parameters. 3) TF-Sapiens: TranscriptFormer-Sapiens trained on 57 million human cells, with 368 million parameters.

In addition to TranscriptFormer, we benchmarked against another cross-species model, namely, Universal Cell Embedding (UCE) (Rosen et al., 2023), which is trained on 36 million cells spanning eight species and 650 million trainable parameters. Besides cross-species models, we benchmarked against several human-only models including Geneformer (Theodoris et al., 2023) (trained on 30 million cells and 30 million parameters), scGPT (Cui et al., 2024) (trained on 33 million human cells and 53 million parameters) and AIDO-a BERT-style encoder-only architecture trained on 50 million cells with four different model sizes, including 3M and 100M (100 million parameters) models used in this study.

We chose the six human tissues from Tabula Sapiens 2 to compare the performance of these models for cell type classification. In Figure 2, we demonstrate that both CytoType and ESM-CE perform comparably and in some cases beat large-scale foundation models in cell type classification. Going forward we define delta as the difference in performance between the lightweight model(s) proposed and the best performing foundation model. Performance is measured using F1-score and a negative value of delta signifies a lower F1-score than the best performing foundation model.

In addition to delta values which are agnostic to which lightweight model is actually the best performing, we measured model-specific delta values (Table 1). For human tissues (Tabula Sapiens 2 dataset), the average delta for CytoType-ESM-Max was −0.053 and the average delta for CytoType-HVG-Max was −0.036. This demonstrates that lightweight models can closely approximate foundation model classification F1 scores on human-derived data. This relatively narrow margin underscores the power of simple baselines. Notably, the simpler ESM-CE variant, which represents a linear layer on top of the average of ESM-2 embeddings for transcribed genes per cell, also performs fairly competitively against computationally expensive foundation models, and in some cases even beats them (Figure 2). Across human tissues, the average delta for ESM-CE was −0.064 for human data.

**Table 1:** Average delta between each lightweight model and the best-performing foundation model across each study. Studies include Tabula Sapiens: 6 human tissues; Cell Atlas: 5 species and multiple tissues per species; Spermatogenesis: 3 species, one tissue each. Sample-weighted averages treat each tissue equally; species-weighted averages (shown for Cell Atlas only) first average within species, then across species. The final column, denoted as “Average delta” represents the difference between the best lightweight model and the best foundation model, averaged across each study. In “Overall Sample” we treat each of the 32 tissue samples from different studies equally and take an average; In “Overall Study” we compute means within each study (using species-weighted averaging for Cell Atlas), then average across the three studies. Closer values to zero indicate closer performance to foundation models.

| Study | CytoType-ESM-Max<br>(average delta) | CytoType-ESM-HVG<br>(average delta) | CytoType-Random-Max<br>(average delta) | ESM-CE<br>(average delta) | Average delta |
| --- | --- | --- | --- | --- | --- |
| Tabula Sapiens | −0.053 | −0.036 | −0.227 | −0.064 | −0.032 |
| Cell Atlas (sample averaging) | −0.083 | −0.098 | −0.234 | −0.066 | −0.046 |
| Cell Atlas (species averaging) | −0.110 | −0.155 | −0.211 | −0.101 | −0.082 |
| Spermatogenesis | −0.019 | −0.058 | −0.062 | −0.013 | −0.011 |
| Overall Sample | −0.071 | −0.083 | −0.217 | −0.061 | −0.040 |
| Overall Study | −0.061 | −0.083 | −0.167 | −0.059 | −0.041 |

Besides benchmarking against human-only scRNA foundation models, we tested the performance of CytoType and ESM-CE against TF and UCE, which are currently the only two models with cross-species training (Figure 3). In particular, we used data from a recent study of spermatogenesis by Murat et al. (2023). Spermatogenesis is the multi-step process by which sperm are produced in the testes. It begins with spermatocytes, which undergo meiosis to generate haploid germ cells. These give rise to early spermatids, round haploid cells that are transcriptionally active and often express newly evolved genes. As development proceeds, later spermatids undergo dramatic remodeling, compacting their DNA, forming the flagellum, and shedding excess cytoplasm to become mature sperm cells. Surrounding somatic cells, particularly Sertoli cells, provide structural support, nutrients, and signaling cues. Together, this coordinated system ensures continuous sperm production and represents one of the most rapidly evolving biological processes in mammals.

**Figure 3:**
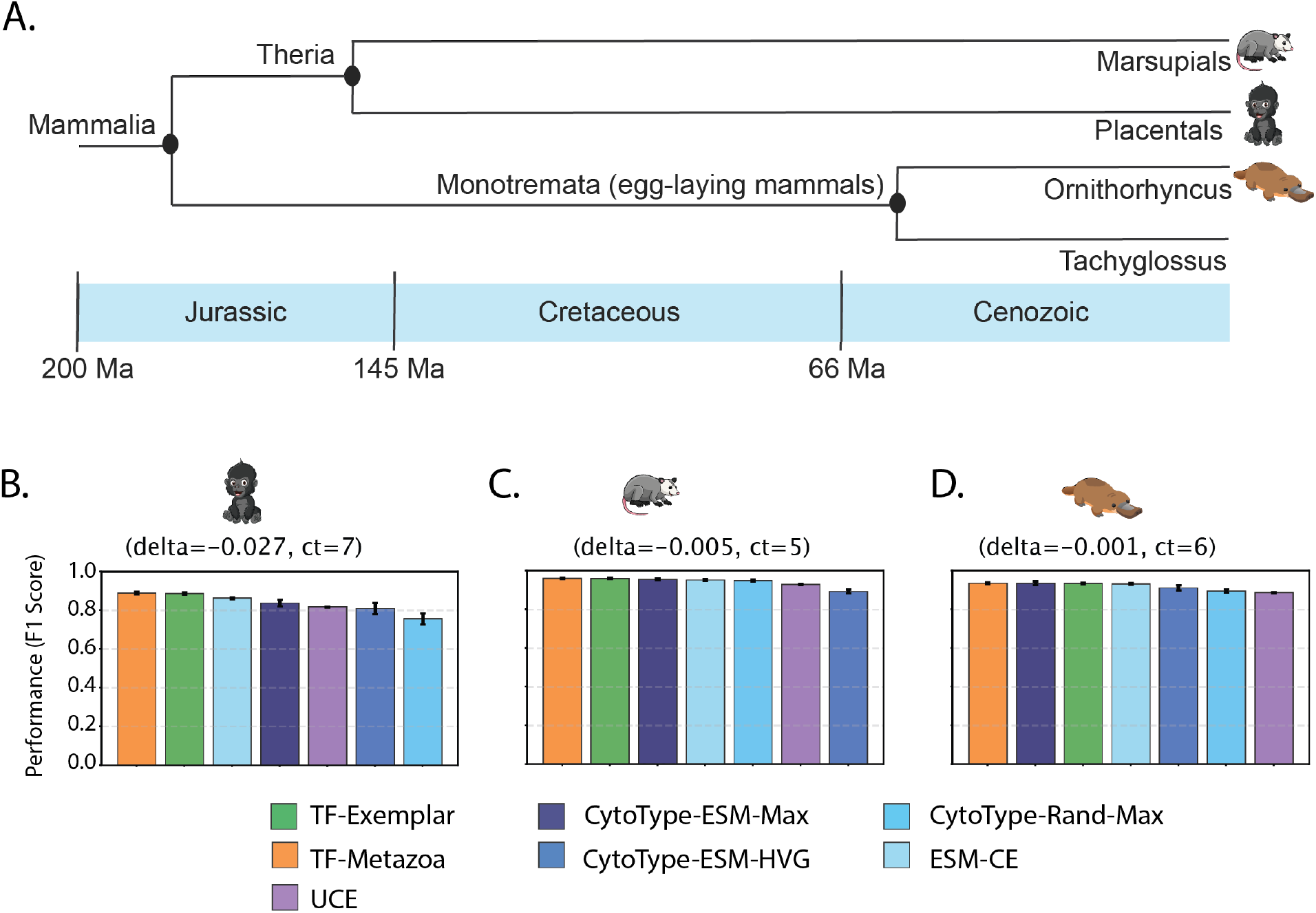
Cross-species benchmarking on a single-tissue dataset. A) Phylogenetic tree of the mammalian clades that each species (B) gorilla, (C) platypus, and (D) opossum belongs to, spanning close 200 Million years of evolutionary distance (tree derived from Whitney, 2025) covering Cenozoic, Cretaceous and Jurassic periods. B-D) Macro F1 scores for various models. The difference between the best performing light-weight model and the best performing foundation model is denoted as delta. The number of cell types in each dataset is represented by “ct”. The models are sorted by their mean F1 score, from highest to lowest.

We compared these models in their ability to classify cell types from gorilla (eutherian), short-tailed opossum (marsupial), and platypus (monotreme), which together represent all three extant mammalian lineages with divergences tracing back to the Mesozoic era and extending into the Cenozoic era, covering more than 145 million years (Whitney, 2025) of evolutionary history (Figure 3). ESM-CE achieves a −0.013 average delta across these species, with Gorilla at −0.027, Opossum at −0.008, and Platypus at just −0.004. CytoType-ESM-Max shows a slightly larger average delta of −0.019 (Figure 3).

Besides spermatogenesis, we benchmarked CytoType and ESM-CE against five other species (Frog, Zebrafish, Mouse Lemur, Coral, and Lamprey), collectively forming the Cell Atlas datasets (Figure 4). Only for Zebrafish and Mouse Lemur, we have data from multiple tissues. The Cell Atlas datasets, encompassing evolutionarily diverse species with diverse samples, present more variable results. CytoType-ESM-Max shows an average delta of −0.11. ESM-CE performs comparably with a −0.10 average delta. As we will discuss, the parameter cost to close performance gaps remains consistently high: approximately 456 million additional parameters on average, which represents 4 orders of magnitude more trainable parameters than CytoType and 5 orders of magnitude more than ESM-CE.

**Figure 4:**
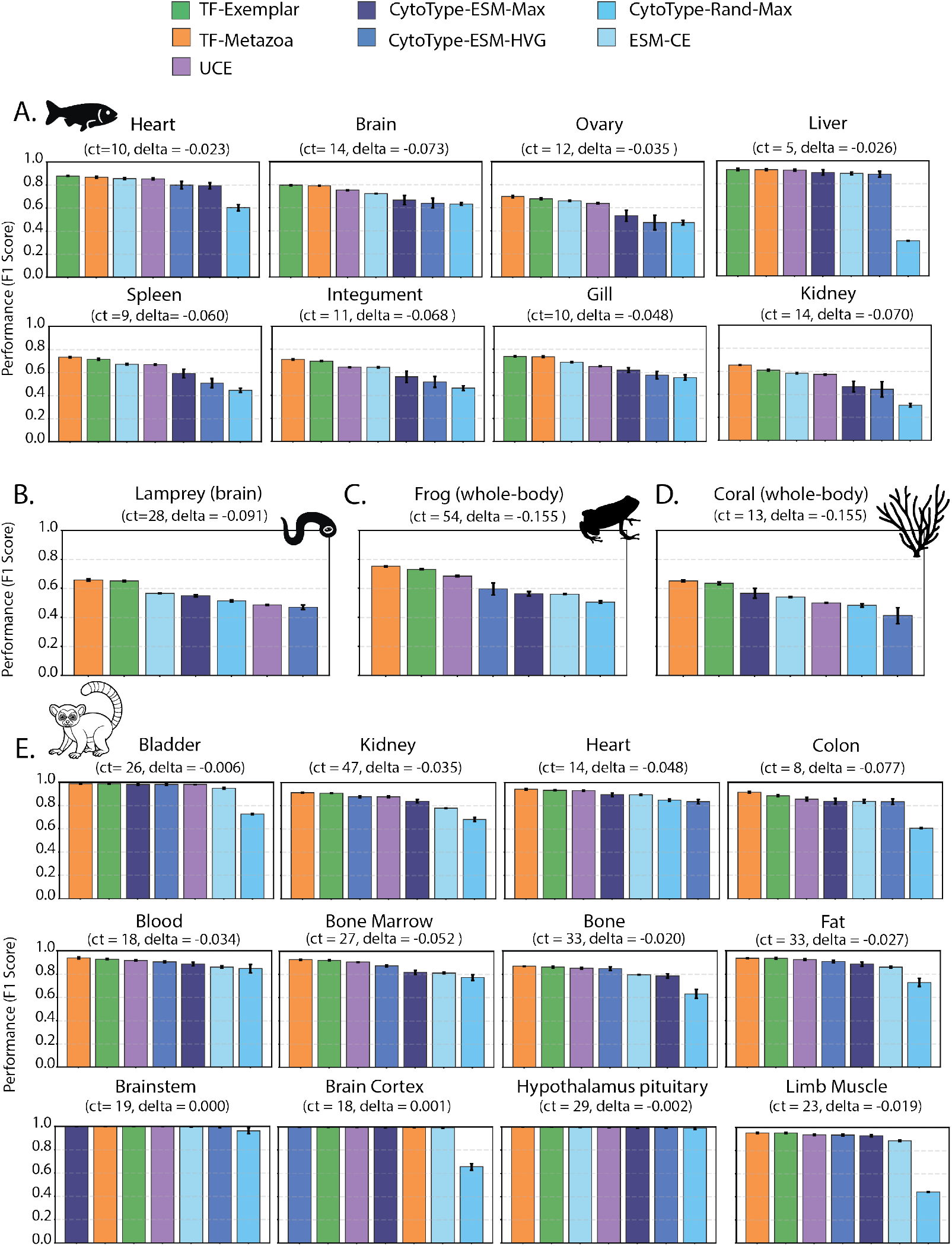
Model F1 scores across the Cell Atlas dataset consisting of five species. A) Zebrafish (8 tissues), B) Lamprey (brain tissue), C) Frog (whole body), D) Coral (whole body), and E) Mouse Lemur (12 tissues). Three cross-species foundation models (UCE, TF-Metazoa, TF-Exemplar) are compared to different variants of CytoType and ESM-CE. The difference between the best performing light-weight model and the best performing foundation model is denoted as delta. The err2o0r bars represent the standard deviation on the five folds. The number of cell types in each dataset is represented by “ct”.

Table 1 summarizes the average deltas between lightweight and foundation models across all datasets. Across all 32 samples (“Overall Sample” in Table 1), the best lightweight model trailed the best foundation model by only delta = −0.04, with ESM-CE achieving the smallest individual gap (delta = −0.061) followed by CytoType-ESM-Max (delta = −0.071) and CytoType-ESM-HVG (delta = −0.083). Study-weighted averaging (“Overall Study”), which gives equal weight to each of the three studies regardless of the number of tissues, yielded similar results (delta = −0.041). Notably, Spermatogenesis showed the smallest performance gaps (delta = −0.011), while Cell Atlas exhibited the largest (delta = −0.082).

### 2.2 ESM embeddings decrease performance gap by three fold compared to random embeddings

Both TranscriptFormer and UCE leverage ESM-2 embeddings as input to enable a universal representation of genes across many species. However, it has not been shown to what extent the ESM-2 embeddings contribute to cell type classification performance. To quantify the contribution of pre-trained ESM-2 embeddings to cell type classification, we compared CytoType-ESM-Max (using ESM-2 embeddings) against CytoType-Random-Max (random gene embeddings) across 32 samples spanning 9 species. CytoType-ESM-Max demonstrated substantial and significantly better performance over the random embedding baseline (Mann-Whitney U = 170, p = 2 *×* 10^−6^), achieving an average delta to the best foundation model of only −0.071 compared to −0.217 for random embedding, which is a three-fold improvement. Notably, every single sample showed better performance with ESM-2 embeddings than random embeddings, with all 32 data points falling above the diagonal in paired comparisons (Figure 5). This performance gain was consistent across all species examined (Figure 5).

**Figure 5:**
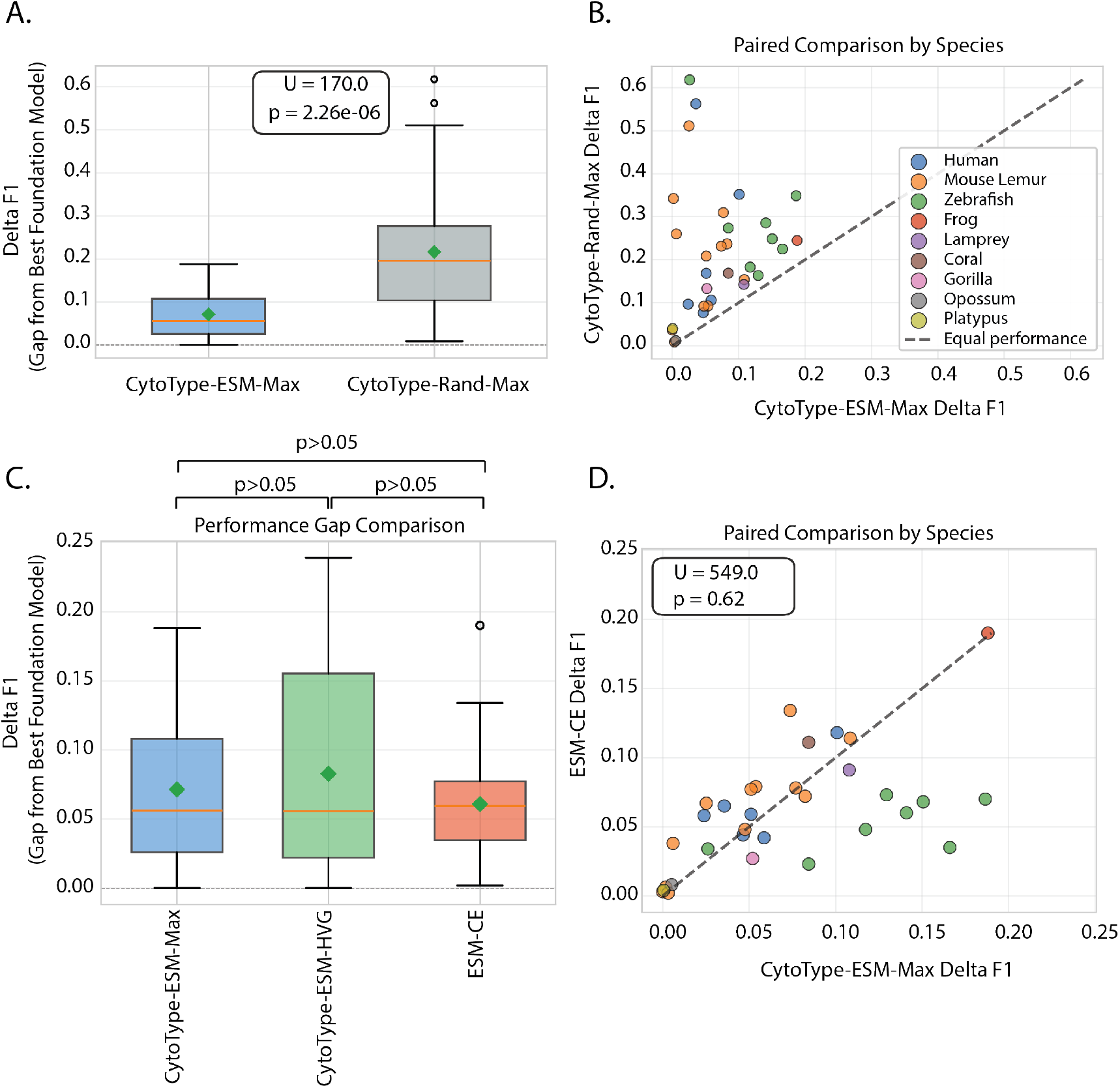
Performance gap analysis comparing ESM-2 embeddings to random embeddings. (A) Performance gap measured as the difference or delta between CytoType variant and the best foundation model: CytoType-ESM-Max (blue) and CytoType-Rand-Max (gray). CytoType-ESM-Max achieves three fold lower delta F1 (mean = 0.071, median = 0.056) compared to random embeddings (mean = 0.217, median = 0.196). Mann-Whitney U test confirms significant difference (U = 170.0, p = 2.26 × 10^−6^). Boxes show interquartile range, whiskers extend to 1.5 × IQR, green diamonds indicate mean values. (B) Paired comparison across 32 samples spanning 9 species. Each point represents one tissue-species combination, colored by species. Points above the diagonal indicate random baseline performs worse than CytoType-ESM-Max. (C) CytoType-ESM-Max (mean = 0.071, median = 0.056), CytoType-ESM-HVG (mean = 0.083, median = 0.056) and ESM-CE (mean = 0.061, median = 0.060) delta F1 compared in a pairwise fashion. No significant difference exists across any of the paired Mann-Whitney U tests performed with Bonferroni correction. (D) Paired comparison across ESM-CE and CytoType-ESM-Max performance gaps, color-coded based on the legend shown in panel B.

We evaluated the two CytoType variants as well as ESM-CE in their delta F1 compared to the best foundation model. ESM-CE achieved the lowest average delta (average delta = −0.061), followed by CytoType-ESM-Max (average delta = −0.071) and CytoType-ESM-HVG (average delta = −0.083), representing only 6-8 point reduction from state-of-the-art foundation models in cell type classification accuracy (Figure 5). However, pairwise statistical testing using Mann-Whitney U tests revealed no significant differences among any variant pairs. ESM-CE demonstrated advantages on non-mammalian species (particularly Zebrafish, average delta = −0.051), while CytoType-ESM-HVG excelled on mammalian tissues (Human: average delta = −0.036, Mouse Lemur: average delta = −0.032), and CytoType-ESM-Max provided balanced performance across taxonomic groups. No single lightweight model dominates across all taxonomic groups; instead, performance differences among lightweight variants are small and context-dependent.

### 2.3 ESM-CE and CytoType variants have comparable cell type classification performance while orders of magnitude smaller in computational complexity compared to foundation models

Foundation models for single-cell transcriptomics, such as UCE, scGPT, and the TF family, differ by several orders of magnitude in number of parameters compared to lightweight methods like CytoType and ESM-CE. The largest model in our evaluation, UCE, comprises approximately 650 million trainable parameters across 33 layers and was pretrained on more than 36 million cells from over 300 datasets for 40 days on 24 A100 80 GB GPUs. The TF-Exemplar and TF-Metazoa models contain roughly 542 million and 444 million parameters, respectively. Even smaller foundation models, such as scGPT (approximately 53 million parameters), still operate at scales several thousand times greater than the parameter budgets of our linear baselines.

These parameter counts for UCE and TF family of models do not include the parameter counts contributed by ESM-2 (since they are not trainable parameters in these models). Because not all foundation models contain ESM-2 embeddings as input, we will restrict the comparison to those that do (Figure 6). For the cross-species classification task, CytoType’s trainable parameter count is given by N cell types *×* N genes (2048) + linear layer embedding dim (2560) + bias, corresponding to 16,897 parameters in gorilla, 14,849 in opossum, and 12,801 in platypus. This corresponds to a reduction of more than 4 orders of magnitude in parameter count relative to even the smallest transformer model evaluated. ESM-CE performs comparably while requiring only 2560 trainable parameters, which is 5 orders of magnitude fewer trainable parameters. These results demonstrate that parameter-efficient baselines remain powerful tools for cell type classification, and that more challenging evaluation tasks are required for the evaluation of single-cell transcriptomics foundation models.

**Figure 6:**
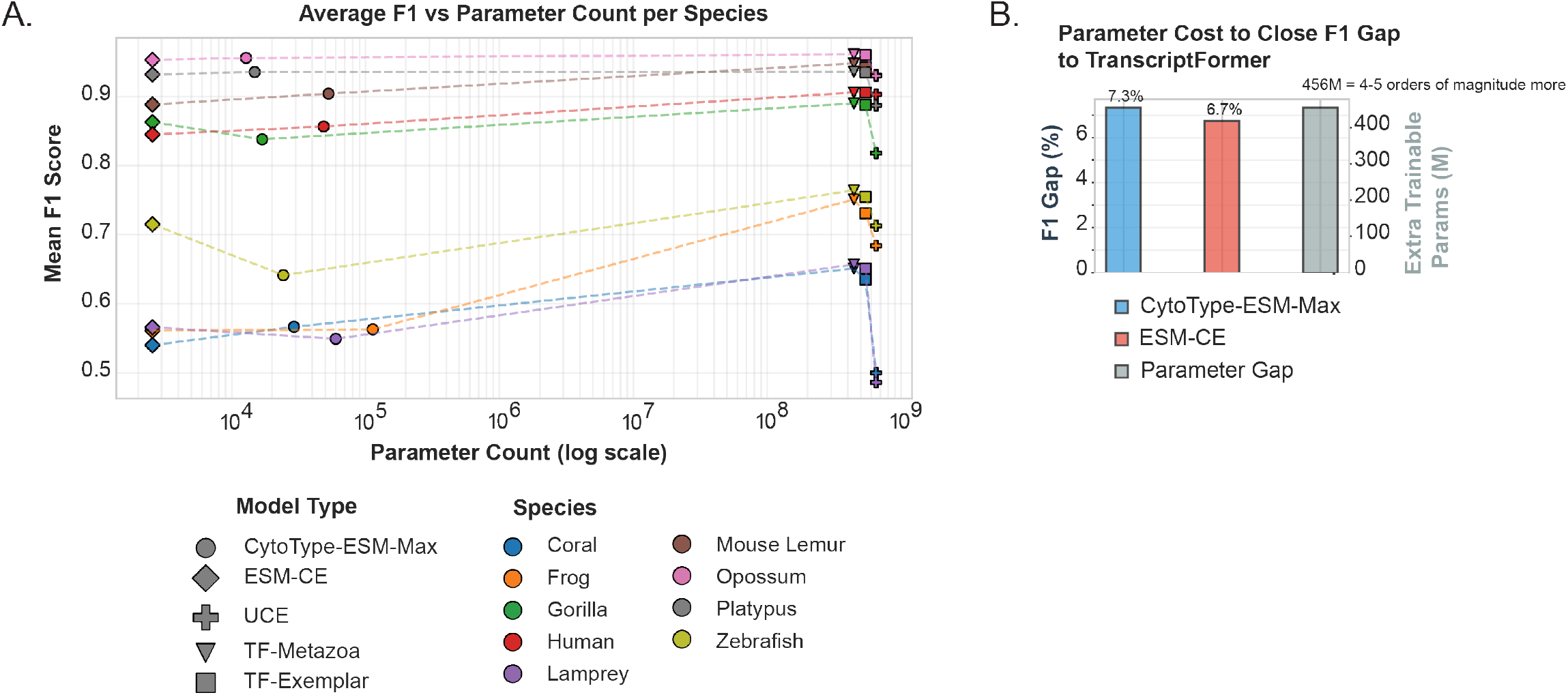
Parameter efficiency comparison. A) Number of parameters (Log10) per model and F1 scores averaged across each species. B) F1 Gap percentage (delta × 100%) versus the parameter gap averaged across species. The difference between either CytoType-ESM-Max or ESM-CE and the best performing foundation model is denoted as delta. The extra parameters is the difference between the number of parameters used in the best foundation model versus either CytoType-ESM-Max or ESM-CE.

**Figure 7:**
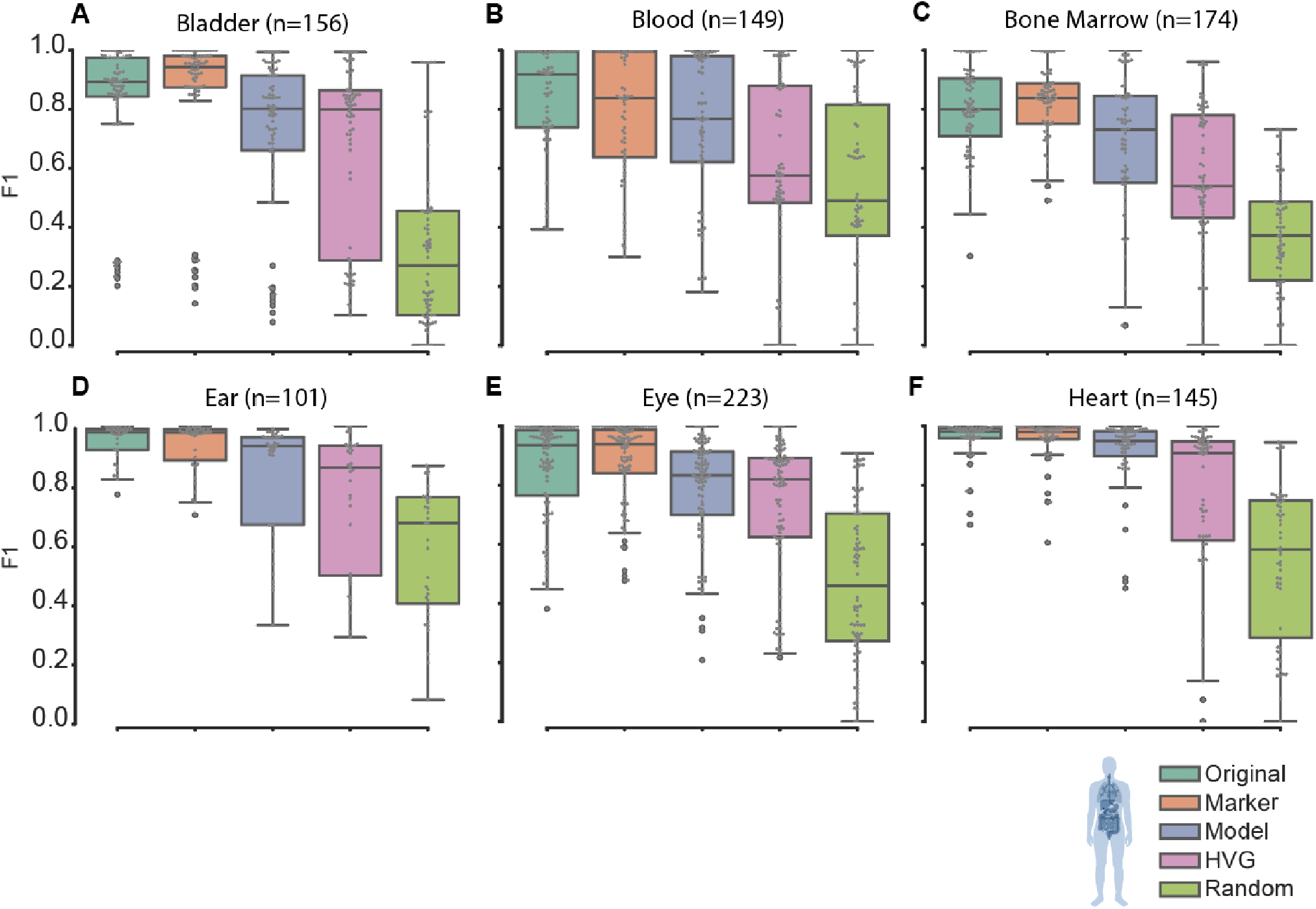
Testing the biological interpretability of the model’s output weights for cell type classification tasks across 6 human tissues. Model-selected genes based on weights (“Model”) are compared to standard marker-gene selection using Scanpy (“Marker”), as well as highly variable genes (“HVG”), and randomly selected genes (“Random”). All these gene selection conditions use the same number of genes (denoted as n), except for “Original” which corresponds to no gene selection. A-F) aggregated F1 scores for several human tissues. G) F1 scores for cell types in heart tissue.

It is important to note that these foundation models were trained across multiple species and tissues, and thus have cell type label transfer capabilities (or out-of-distribution capabilities). On the other hand, CytoType variants and ESM-CE were trained on a specific tissue or species. This leads us to conclude that for specific tasks that do not need generalization across species and tissues, linear models with fewer parameters could perform comparably to foundation models that are computationally expensive and more challenging to train.

### 2.4 CytoType weights are interpretable

Unlike single-cell foundation models whose weights are difficult to interpret, CytoType output is a matrix of cell type specific gene weights. To interpret the output gene weight matrix, we compute a *z*-score based on the distribution of weights for each gene across different cell types for a given tissue (see Materials and Methods). Genes that have *z*-scores above a specified threshold are selected as cell-type specific genes. For each cell type, the genes are ranked by their *z* scores and the top 10 genes per cell type are selected as putative cell type-specific markers (see Materials and Methods). The candidate top genes are collected per tissue and used in cell type classification tasks against other gene-selection methodologies: highly variable genes (“hvg”), marker gene selection based on Scanpy’s differential expression ranking (“marker”), randomly selected genes (“random”), and no gene selection (“original”). In all cases except “original” where we use all available genes, the same number of genes are used to assess the biological interpretability of model weights.

It is important to note that these model-selected genes are not expected to outperform full gene sets or marker genes selected using standard differential expression methods, which have access to quantitative count information. Rather, this analysis is designed to test whether CytoType’s learned weights—despite being trained without access to expression counts—encode biologically meaningful information that can support downstream cell type discrimination.

Thus, we evaluated the classification performance across different gene selection strategies using the reduced collection of genes, going from greater than 2000 genes to less than 200. We evaluated per-cell type F1 scores using a stratified *k* fold cross-validation. For each tissue, we computed PCA on the log-normalized count data, and used a *k*-nearest neighbors (*k*-NN) classifier to predict cell type labels. Per-class (i.e., per-cell type) F1 scores are computed for each fold (see Methods for further details).

Model selected genes consistently outperformed both “HVG” and “Random” baselines across all tissues (all FDR *<* 0.05), indicating that model weights despite being trained without count information contain fairly small sets of genes that are indicative of cell types (Figure 7). As expected, model-selected genes did not outperform full sets of genes or marker genes selected through standard marker selection methodology which relies on count information. Without the count information, however, the genes selected by the model perform fairly well. For example, the median F1 scores for model-selected genes were 0.80 (bladder, 0.72 *±* 0.26), 0.77 (blood, 0.74 *±* 0.25), 0.73 (bone marrow, 0.68 *±* 0.23), 0.93 (ear, 0.83 *±* 0.20), 0.83 (eye, 0.78 *±* 0.18), and 0.95 (heart, 0.91 *±* 0.12). We report the median as certain cell-type specific F1 scores are outliers due to the small number of cells associated with the cell type. For example, in the case of heart tissue, the model selected genes have the worst performance for monocytes, which with 111 representative cells only marginally passes the threshold number of observations that we impose, which is 100 cells. In general, we show that CytoType’s learned weights enable cell type classification.

## 3 Discussion & Limitations

Our findings show that small, interpretable models can perform competitively with large-scale foundation models for single-cell classification. Both CytoType and ESM-CE achieve high F1 scores across tissues and species despite containing only thousands of parameters—four or five orders of magnitude fewer than state-of-the-art transformer-based architectures. This extreme parameter efficiency highlights that strong cell type classification performance does not require billion-parameter networks or large-scale pretraining.

For well-defined cell type classification tasks within a specific species and tissue context where cross-species generalization is not required the proposed lightweight models achieve comparable performance to foundation models with orders of magnitude fewer parameters. The strong performance of such lightweight models parallels findings from recent systematic bench-marking of single-cell foundation models (Ahlmann-Eltze et al., 2025) (`scGPT`, `scFoundation`, `Geneformer`, `scBERT`, `UCE`) and specialized perturbation model against deliberately simple baseline models for predicting transcriptomic responses to genetic perturbations. Together, these results suggest that scaling model size alone is not sufficient to achieve better biological understanding or predictive accuracy for certain biological tasks. It also suggests that cell type classification, which is often a critical evaluation task used in this field, is a relatively simple computational task that does not scale with parameters used.

Despite these strengths, our study has several limitations. First, CytoType and ESM-CE are task-specific models designed exclusively for cell type classification and do not provide the broad representational reuse, generative capabilities, or multi-task flexibility offered by large foundation models. Second, the lightweight models are trained and evaluated within individual species or tissue contexts and are not designed to support cross-species or crosstissue label transfer, a setting in which cross-species foundation models are expected to excel. As a result, while lightweight models offer strong performance and interpretability for targeted classification tasks, large foundation models remain better suited for more general or exploratory single-cell analyses.

## 4 Conclusion

In this work, we introduce CytoType and ESM-CE, two lightweight models that leverage pretrained ESM-2 protein embeddings to classify cell types using only binary gene expression information during training, achieving performance comparable to large single-cell foundation models while using four to five orders of magnitude fewer trainable parameters. Across nine species and more than thirty tissues, these results demonstrate that much of the signal required for accurate cell type classification is captured by the presence or absence of transcripts rather than their quantitative expression levels. Pretrained ESM-2 embeddings substantially reduce the performance gap to state-of-the-art foundation models by approximately three fold compared to random embeddings. Notably, even random embeddings perform unexpectedly well. In addition to strong performance, CytoType provides biologically interpretable, genespecific weights that identify cell-type–associated gene sets despite being trained without access to expression counts. Together, these findings show that for targeted cell type classification within a fixed species or tissue, simple and interpretable models can recover much of the accuracy of large foundation models at a fraction of the computational cost.

## 5 Methods

### 5.1 CytoType and ESM-CE

To recap, our goal is to predict cell types given single cell transcriptomic counts. Our datasets spans 9 species and 30+ tissues. Each species has a different gene vocabulary. The first step was to perform gene selection and aggregate the gene embeddings of the selected genes to represent each cell. This approach enables us to (a) use a lighter weight model, and (b) ensure cells from species with different gene vocabulary lengths are trained on the same dimensionality. We used ESM-2 embeddings generated for TranscriptFormer. Briefly, for each gene in each species, its associated proteins sequence(s) is obtained and these sequences are embedded using ESM-2 model. In cases where there are multiple proteins associated with a gene the average over all ESM-2 protein embeddings is used.

This representation is then used as input to a model that uses a cross-entropy loss function to classify cell type. The cross-entropy loss function is defined as:

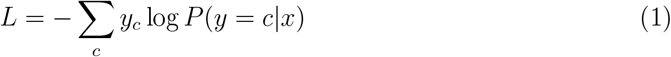

where *y*_*c*_ is the one-hot encoded true label.

We implemented several lightweight models, which differ in gene selection methodology and training function.

#### Gene Selection Methodology

We follow a similar procedure to TranscriptFormer and select the top *k* = 2048 genes for each sample (or the full gene vocabulary length if fewer than 2048 genes) based on the following criteria. The ESM-2 embeddings of the selected genes for a given sample are then linearly combined (averaged) to form a cell representation.

Let us denote the selected gene set by *G*.

- **Most highly expressed genes:** Genes with the highest expression value where the expected expression value across cells is greater than zero
- **Most highly variable genes:** Genes with the highest variability across cells in each (species, tissue) group, determined using Scanpy’s `highly variable genes` function (Seurat flavor)

**ESM-CE** predicts the probability of a cell belonging to a specific cell type based on its averaged ESM-2 gene embedding for the 2048 most expressed genes. For each sample (i.e., a cell), we compute a logit *s*_*c*_ for each candidate cell type *c* using the following formulations. A softmax function is used to estimate probabilities followed by cross entropy loss.

#### CytoType

Here, we learn two weight vectors, *v* ∈ ℝ^2560^ and 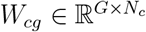, where *c* is the number of cell types. The *W*_*cg*_ matrix is constrained to be non-negative. The model not only learns the importance of genes in a cell type, but also how to read the genes via *v*. The formulation is as follows:

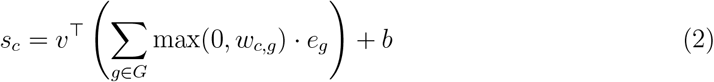

where *G* is the selected gene set using the methodologies above, *e*_*g*_ is the ESM-2 embedding for gene *g* and *b* is a bias term. To obtain the probability distribution over cell-types, we apply the softmax functions over the logits.

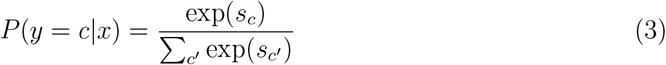

The model is trained using the cross-entropy loss, as defined in Equation (1).

Combining the above gene selection methods and models we arrived at the following lightweight models in Table 2:

**Table 2:** Model descriptions.

| Model | Description |
| --- | --- |
| ESM-CE | Maximum expressed genes + ESM2 gene embeddings + logistic regression |
| CytoType-ESM-Max | Maximum expressed genes + ESM2 gene embeddings + CytoType |
| CytoType-ESM-HVG | Most highly variable genes + ESM2 gene embeddings + CytoType |
| CytoType-Random-Max | Maximum expressed genes + embeddings from $\mathcal{N}(0, 1)$ + CytoType |

The models were trained for 5 epochs and results were reported with a 5 fold cross validation.

#### Foundation Baselines

CytoType and ESM-CE were benchmarked against eight foundation models trained on single-cell transcriptomic data: TF-Metazoa, TF-Exemplar, TF-Sapiens, scGPT, GeneFormer, UCE, AIDO-3M, and AIDO-100M. For each foundation model, cell embeddings were extracted from the final transformer layer and normalized to zero mean and unit variance. A logistic regression classifier was then trained on these normalized embeddings for cell type classification.

#### Performance Comparison Analysis

To quantify the performance gap between lightweight models (CytoType-ESM-Max, CytoType-ESM-HVG, CytoType-Random-Max, and ESM-CE) and foundation models, we calculated delta values, where delta = Lightweight Model F1 − Best Foundation Model F1. Negative values indicate that foundation models outperform lightweight models. For each sample, we identified the best foundation model by selecting the maximum F1 score among TF-Metazoa, TF-Exemplar, UCE, TF-Sapiens, scGPT, Geneformer, scVI, aido-cell-100m, and aido-cell-3m. We computed both model-specific delta values (the difference between each lightweight model and the best foundation model) and the difference between the best lightweight model and the best foundation model.

Summary statistics were computed at two levels. At the dataset level, we used species-weighted averaging for Cell Atlas, where delta scores are first averaged within each species then averaged across species to ensure equal contribution regardless of the number of tissues sampled per species. At the overall level, we computed: (1) sample-weighted averages (“Overall Sample” in Table 1), treating each of the 32 samples equally, and (2) study-weighted averages (“Overall Study” in Table 1), where we first computed the species-weighted mean (applies only to Cell Atlas study) within each of the three studies, then averaged across studies.

### 5.2 CytoType’s Weight Interpretability Analysis

Single-cell RNA sequencing data were stored and analyzed in AnnData objects (v0.10.0) using Scanpy (v1.9.8). For each dataset, raw counts were normalized with a log-transformation. Dimensionality reduction was performed by principal component analysis (PCA) using up to 40 components, constrained by the number of cells. Neighborhood graphs were computed on the PCA representation with 10 nearest neighbors. Uniform Manifold Approximation and Projection (UMAP) embeddings were generated for visualization.

#### 5.2.1 Model-Derived Weights and Gene Selection

Cell-type–specific gene weight matrices were loaded from pretrained models (`weights.pt`) which we refer to as *W*_*cg*_, where *c* is the number of cell types and *g* is the number of genes. The entry *W*_*ij*_ represents the contribution of gene *j* to cell type *i*.

To quantify the cell type-specificity of each gene, we compute a *z*-score for gene *j* in cell type *i* relative to its distribution across all other cell types. Define the mean and standard deviation of gene *j*’s weights across all cell types except *i* as:

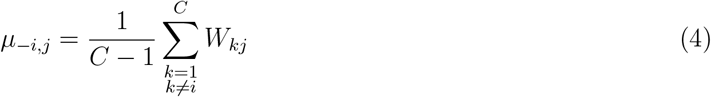

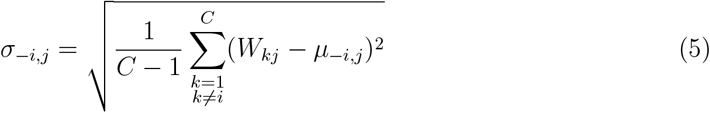

The specificity *z*-score for gene *j* in cell type *i* is then defined as:

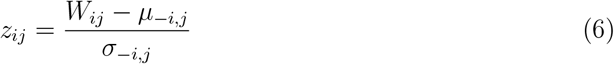

with the convention that if *σ*_−*i,j*_ = 0 then *z*_*ij*_ = 0.

A gene is considered a specific marker for cell type *i* if:

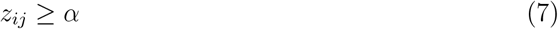

where *α* is a predetermined threshold (e.g., *α* = 3). For each cell type, the genes are ranked by their *z*_*ij*_ values and the top *n* genes are selected as cell type-specific markers. Genes with *z*_*ij*_ exceeding a chosen threshold *α* are then considered important to a cell type’s identity *i*. Genes with *z*-scores above 3.0 were retained, and up to the top 10 genes per cell type were selected. This yielded sets of model-selected genes, which were compared against three alternative gene selection strategies:

1. **Marker genes:** computed by differential expression ranking (`sc.tl.rank genes groups`) with a *t*-test, and sub-sampled to match the number of model-selected genes.
2. **Highly variable genes (HVGs):** determined using Scanpy’s `highly variable genes` function (Seurat flavor) with 2000 top HVGs, followed by random sub-sampling.
3. **Random genes:** sampled uniformly at random from all expressed genes.

#### 5.2.2 Comparative Embedding and Visualization

For each gene set (original, model-selected, marker, HVG, random), AnnData objects were subset to the chosen features and reprocessed through PCA, neighbor graph construction, Leiden clustering, and UMAP projection. UMAPs were visualized with Scanpy’s plotting functions.

#### 5.2.3 Evaluation of Per-Cell Type F1 Scores Across Different Gene Selection Methodologies

Classification performance was assessed by training *k*-nearest neighbors (*k*-NN) classifiers (*k* = 5) on the PCA coordinates in a stratified 5-fold cross-validation scheme. F1 scores were computed per cell type for each fold. Raw per-cell type F1 scores were aggregated across folds and experimental conditions (Original, Marker, Model, HVG, Random).

#### 5.2.4 Visualization of Performance Metrics

Performance comparisons across conditions were visualized using grouped and aggregated boxplots. Distributions of F1 scores were displayed per cell type, ordered by mean F1 score, and across all cell types for each condition. Visualizations were generated using Seaborn (v0.13.2) and Matplotlib (v3.9.2). Analyses were performed independently for each human tissue using corresponding pretrained weight matrices.

#### 5.2.5 Statistical Testing

We used Wilcoxon signed-rank tests to compare the paired distributions of F1 scores between the Model condition and other gene selection conditions (HVG, Marker, Random, Original) within each dataset. One-sided alternatives were specified based on expectations:

- For HVG and Random baselines, we tested whether Model achieved higher F1 scores (Model performance better than baseline).
- For Marker and Original baselines, we tested whether Model achieved lower F1 scores (Model performance not better than baseline).

Raw *p*-values were adjusted for multiple comparisons using the Benjamini–Hochberg false discovery rate (FDR) procedure within each dataset.

## 6 Code Availability

Training code as well as weight analyses code and notebooks are provided in this repository: https://github.com/czi-ai/CytoType

## 7 Data Availability

All datasets presented in this manuscript have been previously provided and described as part of our earlier work on TranscriptFormer (Pearce et al., 2025). Here is a brief recap:

- Tabula Sapiens 2.0 (Quake et al., 2025) is a human scRNA-seq reference cell atlas originating from Tabula Sapiens Consortium via CZ CELLxGENE.
- Spermatogenesis dataset is a single-tissue (testes), multi-species snRNA-seq dataset from several lineages of mammals and birds (Murat et al., 2023).
- Tabula Microcebus data is a scRNA-seq cell atlas for mouse lemur (Microcebus murinus), obtained from the Tabula Microcebus Consortium (Consortium et al., 2021).
- Tropical clawed frog cell atlas is a scRNA-seq developmental cell atlas for X. tropicalis covering embryonic stages (Briggs et al., 2018).
- Zebrafish dataset is a scRNA-seq developmental cell atlas for D. rerio (Jiang et al., 2021). The dataset was subset to the adult stage and split by tissue.
- Stony coral scRNA-seq atlas (Stylophora pistillata) (Levy et al., 2021) was subset to include adult stage only.
- Sea lamprey brain dataset is a snRNA-seq dataset from Petromyzon marinus (Lamanna et al., 2023).

## 8 Acknowledgements

We would like to thank all members of the TranscriptFormer team including Sara Simmonds, Giovanni Palla, Ana-Maria Istrate, Alexander Tarashansky, Benjamin Nelson, Omar Valenzuela, and Donghui Li. We also thank Jakub Tomczak for helpful suggestions.

